# A combined proteomics and Mendelian randomization approach to investigate the effects of aspirin-targeted proteins on colorectal cancer

**DOI:** 10.1101/2020.08.14.239871

**Authors:** Aayah Nounu, Alexander Greenhough, Kate J Heesom, Rebecca C Richmond, Jie Zheng, Stephanie J Weinstein, Demetrius Albanes, John A Baron, John L Hopper, Jane C Figueiredo, Polly A Newcomb, Noralane M Lindor, Graham Casey, Elizabeth A Platz, Loïc Le Marchand, Cornelia M Ulrich, Christopher I Li, Fränzel JB van Duijnhoven, Andrea Gsur, Peter T Campbell, Víctor Moreno, Pavel Vodicka, Ludmila Vodickova, Hermann Brenner, Jenny Chang-Claude, Michael Hoffmeister, Lori C Sakoda, Martha L Slattery, Robert E Schoen, Marc J Gunter, Sergi Castellví-Bel, Hyeong Rok Kim, Sun-Seog Kweon, Andrew T Chan, Li Li, Wei Zheng, D Timothy Bishop, Daniel D Buchanan, Graham G Giles, Stephen B Gruber, Gad Rennert, Zsofia K Stadler, Tabitha A Harrison, Yi Lin, Temitope O Keku, Michael O Woods, Clemens Schafmayer, Bethany Van Guelpen, Steven J Gallinger, Heather Hampel, Sonja I Berndt, Paul D P Pharoah, Annika Lindblom, Alicja Wolk, Anna H Wu, Emily White, Ulrike Peters, David A Drew, Dominique Scherer, Justo Lorenzo Bermejo, Ann C Williams, Caroline L Relton

**Affiliations:** Medical Research Council (MRC) Integrative Epidemiology Unit, Bristol Medical School, University of Bristol, Bristol, BS8 2BN, UK; School of Cellular and Molecular Medicine, University of Bristol, Bristol, BS8 1TD, UK; Centre for Research in Biosciences, The Faculty of Health and Applied Sciences, The University of the West of England, Bristol, BS16 1QY, UK; Proteomics Facility, Faculty of Life Sciences, University of Bristol, Bristol, UK; Division of Cancer Epidemiology and Genetics, National Cancer Institute, National Institutes of Health, Bethesda, Maryland, USA; Department of Medicine, University of North Carolina School of Medicine, Chapel Hill, North Carolina, USA; Centre for Epidemiology and Biostatistics, Melbourne School of Population and Global Health, The University of Melbourne, Melbourne, Victoria, Australia; Department of Epidemiology, School of Public Health and Institute of Health and Environment, Seoul National University, Seoul, South Korea; Department of Medicine, Samuel Oschin Comprehensive Cancer Institute, Cedars-Sinai Medical Center, Los Angeles, CA, USA; Department of Preventive Medicine, Keck School of Medicine, University of Southern California, Los Angeles, California, USA; Public Health Sciences Division, Fred Hutchinson Cancer Research Center, Seattle, Washington, USA; School of Public Health, University of Washington, Seattle, Washington, USA; Department of Health Science Research, Mayo Clinic, Scottsdale, Arizona, USA; Center for Public Health Genomics, University of Virginia, Charlottesville, Virginia, USA; Department of Epidemiology, Johns Hopkins Bloomberg School of Public Health, Baltimore, Maryland, USA; University of Hawaii Cancer Center, Honolulu, Hawaii, USA; Huntsman Cancer Institute and Department of Population Health Sciences, University of Utah, Salt Lake City, Utah, USA; Division of Human Nutrition and Health, Wageningen University & Research, Wageningen, The Netherlands; Institute of Cancer Research, Department of Medicine I, Medical University Vienna, Vienna, Austria; Behavioral and Epidemiology Research Group, American Cancer Society, Atlanta, Georgia, USA; Oncology Data Analytics Program, Catalan Institute of Oncology-IDIBELL, L'Hospitalet de Llobregat, Barcelona, Spain; CIBER Epidemiología y Salud Pública (CIBERESP), Madrid, Spain; Department of Clinical Sciences, Faculty of Medicine, University of Barcelona, Barcelona, Spain; ONCOBEL Program, Bellvitge Biomedical Research Institute (IDIBELL), L'Hospitalet de Llobregat, Barcelona, Spain; Department of Molecular Biology of Cancer, Institute of Experimental Medicine of the Czech Academy of Sciences, Prague, Czech Republic; Institute of Biology and Medical Genetics, First Faculty of Medicine, Charles University, Prague, Czech Republic; Faculty of Medicine and Biomedical Center in Pilsen, Charles University, Pilsen, Czech Republic; Division of Clinical Epidemiology and Aging Research, German Cancer Research Center (DKFZ), Heidelberg, Germany; Division of Preventive Oncology, German Cancer Research Center (DKFZ) and National Center for Tumor Diseases (NCT), Heidelberg, Germany; German Cancer Consortium (DKTK), German Cancer Research Center (DKFZ), Heidelberg, Germany; Division of Cancer Epidemiology, German Cancer Research Center (DKFZ), Heidelberg, Germany; University Medical Centre Hamburg-Eppendorf, University Cancer Centre Hamburg (UCCH), Hamburg, Germany; Division of Research, Kaiser Permanente Northern California, Oakland, California, USA; Department of Internal Medicine, University of Utah, Salt Lake City, Utah, USA; Department of Medicine and Epidemiology, University of Pittsburgh Medical Center, Pittsburgh, Pennsylvania, USA; Nutrition and Metabolism Section, International Agency for Research on Cancer, World Health Organization, Lyon, France; Gastroenterology Department, Hospital Clínic, Institut d'Investigacions Biomèdiques August Pi i Sunyer (IDIBAPS), Centro de Investigación Biomédica en Red de Enfermedades Hepáticas y Digestivas (CIBEREHD), University of Barcelona, Barcelona, Spain; Department of Surgery, Chonnam National University Hwasun Hospital and Medical School, Hwasun, Korea; Department of Preventive Medicine, Chonnam National University Medical School, Gwangju, Korea; Jeonnam Regional Cancer Center, Chonnam National University Hwasun Hospital, Hwasun, Korea; Division of Gastroenterology, Massachusetts General Hospital and Harvard Medical School, Boston, Massachusetts, USA; Channing Division of Network Medicine, Brigham and Women's Hospital and Harvard Medical School, Boston, Massachusetts, USA; Clinical and Translational Epidemiology Unit, Massachusetts General Hospital and Harvard Medical School, Boston, Massachusetts, USA; Broad Institute of Harvard and MIT, Cambridge, Massachusetts, USA; Department of Epidemiology, Harvard T.H. Chan School of Public Health, Harvard University, Boston, Massachusetts, USA; Department of Immunology and Infectious Diseases, Harvard T.H. Chan School of Public Health, Harvard University, Boston, Massachusetts, USA; Department of Family Medicine, University of Virginia, Charlottesville, Virginia, USA; Division of Epidemiology, Department of Medicine, Vanderbilt-Ingram Cancer Center, Vanderbilt Epidemiology Center, Vanderbilt University School of Medicine, Nashville, Tennessee, USA; Leeds Institute of Cancer and Pathology, University of Leeds, Leeds, UK; Colorectal Oncogenomics Group, Department of Clinical Pathology, The University of Melbourne, Parkville, Victoria 3010 Australia; University of Melbourne Centre for Cancer Research, Victorian Comprehensive Cancer Centre, Parkville, Victoria 3010 Australia; Genetic Medicine and Family Cancer Clinic, The Royal Melbourne Hospital, Parkville, Victoria, Australia; Cancer Epidemiology Division, Cancer Council Victoria, Melbourne, Victoria, Australia; Precision Medicine, School of Clinical Sciences at Monash Health, Monash University, Clayton, Victoria, Australia; Department of Preventive Medicine & USC Norris Comprehensive Cancer Center, Keck School of Medicine, University of Southern California, Los Angeles, California, USA; Department of Community Medicine and Epidemiology, Lady Davis Carmel Medical Center, Haifa, Israel; Ruth and Bruce Rappaport Faculty of Medicine, Technion-Israel Institute of Technology, Haifa, Israel; Clalit National Cancer Control Center, Haifa, Israel; Department of Medicine, Memorial Sloan Kettering Cancer Center, New York, New York, USA; Center for Gastrointestinal Biology and Disease, University of North Carolina, Chapel Hill, North Carolina, USA; Memorial University of Newfoundland, Discipline of Genetics, St. John's, Canada; Department of General Surgery, University Hospital Rostock, Rostock, Germany; Department of Radiation Sciences, Oncology Unit, Umeå University, Umeå, Sweden; Wallenberg Centre for Molecular Medicine, Umeå University, Umeå, Sweden; Lunenfeld Tanenbaum Research Institute, Mount Sinai Hospital, University of Toronto, Toronto, Ontario, Canada; Division of Human Genetics, Department of Internal Medicine, The Ohio State University Comprehensive Cancer Center, Columbus, Ohio, USA; Department of Public Health and Primary Care, University of Cambridge, Cambridge, UK; Department of Clinical Genetics, Karolinska University Hospital, Stockholm, Sweden; Department of Molecular Medicine and Surgery, Karolinska Institutet, Stockholm, Sweden; Institute of Environmental Medicine, Karolinska Institutet, Stockholm, Sweden; Department of Surgical Sciences, Uppsala University, Uppsala, Sweden; University of Southern California, Preventative Medicine, Los Angeles, California, USA; Department of Epidemiology, University of Washington School of Public Health, Seattle, Washington, USA; Massachusetts General Hospital and Harvard Medical School, Clinical and Translational Epidemiology Unit, Boston, Massachusetts 02114, USA; Institute of Medical Biometry and Informatics, University of Heidelberg, Im Neuenheimer Feld 130.3, Heidelberg, Germany

**Keywords:** aspirin, proteome, Mendelian randomization, colorectal

## Abstract

**Background:** Evidence for aspirin’s chemopreventative properties on colorectal cancer (CRC) is substantial, but its mechanism of action is not well-understood. We combined a proteomic approach with Mendelian randomization (MR) to identify possible new aspirin targets that decrease CRC risk.

**Methods:** Human colorectal adenoma cells (RG/C2) were treated with aspirin (24 hours) and a stable isotope labelling with amino acids in cell culture (SILAC) based proteomics approach identified altered protein expression. Protein quantitative trait loci (pQTLs) from INTERVAL (N=3,301) and expression QTLs (eQTLs) from the eQTLGen Consortium (N=31,684) were used as genetic proxies for protein and mRNA expression levels. Two-sample MR of mRNA/protein expression on CRC risk was performed using eQTL/pQTL data combined with CRC genetic summary data from the Colon Cancer Family Registry (CCFR), Colorectal Transdisciplinary (CORECT), Genetics and Epidemiology of Colorectal Cancer (GECCO) consortia and UK Biobank (55,168 cases and 65,160 controls).

**Results:** Altered expression was detected for 125/5886 proteins. Of these, aspirin decreased MCM6, RRM2 and ARFIP2 expression and MR analysis showed that a standard deviation increase in mRNA/protein expression was associated with increased CRC risk (OR:1.08, 95% CI:1.03-1.13, OR:3.33, 95% CI:2.46-4.50 and OR:1.15, 95% CI:1.02-1.29, respectively).

**Conclusion:** MCM6 and RRM2 are involved in DNA repair whereby reduced expression may lead to increased DNA aberrations and ultimately cancer cell death, whereas ARFIP2 is involved in actin cytoskeletal regulation indicating a possible role in aspirin’s reduction of metastasis.

**Impact:** Our approach has shown how laboratory experiments and population-based approaches can combine to identify aspirin-targeted proteins possibly affecting CRC risk.

## Introduction

Colorectal cancer (CRC) is the fourth most common cancer worldwide (1). Observational studies as well as randomized controlled trials (RCTs) using aspirin for the prevention of vascular events have shown that aspirin use is associated with a decrease in CRC incidence and mortality (2–5). This was primarily thought to be through the acetylation of the cyclooxygenase (COX) enzymes thereby inhibiting their action (6). These enzymes are involved in the COX/prostaglandin E2(PGE_2_) signalling pathway which is frequently upregulated in CRC, driving many of the hallmarks of cancer (7,8).

Evidence for COX-independent mechanisms have also emerged, such as the prevention of NFκB activation, inhibition of the extracellular-signal-regulated kinase (ERK) signalling pathway, cell cycle progression inhibition and possible induction of autophagy (7,9). An aspirin derivative that does not inhibit COX reduced the mean number of aberrant crypt foci (an early lesion in colorectal carcinogenesis) in a mouse model of CRC more than aspirin itself (10). Furthermore, aspirin was able to inhibit proliferation and induce apoptosis in COX-2 negative colon cancer cell lines as well as reducing angiogenesis in 3D assays where COX-inhibitors showed no effect (11–13). Clinically, aspirin has been shown to reduce tumour recurrence in phosphatidylinositol-4,5-bisphosphate 3-kinase catalytic subunit alpha (PIK3CA) mutant cancer whereas rofecoxib (a COX-2 selective inhibitor) showed no effect (14) and has also been shown to improve survival in patients with human leukocyte antigen (HLA) class I antigen expression, regardless of COX-2 expression (15). There is now a significant number of studies that indicate the mechanism behind the action of aspirin on CRC risk is still not fully understood and that multiple mechanisms are involved (16).

In conventional epidemiological studies it is often difficult to determine causality due to limitations of confounding and reverse causation. While RCTs can overcome these limitations, they are generally limited to assessing the causal role of health interventions or pharmaceutical agents on disease outcomes, rather than understanding biological mechanisms. Furthermore, in the context of cancer, RCTs for cancer primary prevention are not always feasible, as they require long-term follow-up for the cancer to develop. Mendelian randomization (MR) is an epidemiological method which applies a similar notion of randomization as in the RCT to evaluate causality. In MR, genetic variants (most commonly single nucleotide polymorphisms (SNPs)) are used to proxy an exposure of interest (17). As genetic variants are randomly assorted at conception, an individual’s genetic makeup is unlikely to be influenced by exposures later on in life, thus reducing the possibility of confounding and reverse causation (18).

More recently, the increase in genome-wide association studies for molecular traits has identified SNPs that are associated with protein and mRNA expression levels, thereby providing protein quantitative trait loci (pQTLs) and expression quantitative trait loci (eQTLs) (19,20), which may be used to investigate the causal mechanism of drug targets on disease risk (21). Such methods can complement laboratory experiments to better understand the mechanism of action of drugs on cancer growth and progression.

Due to evidence showing that aspirin may prevent adenoma formation (22) and adenomas being the precursors of most colorectal cancers (23), we focused on a colorectal adenoma cell line (RG/C2) in this study and identified altered protein expression in relation to aspirin treatment. Findings were then taken forward into an MR analysis to investigate which proteins targeted by aspirin may be causally implicated in reducing risk of CRC incidence, thereby providing insight into alternative mechanisms/pathways for the action of aspirin.

## Methods

### Cell culture experiments

The S/RG/C2 (referred to as RG/C2 henceforth whereby the prefix “S” denotes that they are from a sporadic tumour) (RRID:CVCL_IQ11) colorectal adenoma cell line was derived in the Colorectal Tumour Biology group and is described in detail elsewhere (24). These cells express RG/C2 cells express WT full length *APC* (251) as well as wild type *KRAS* and *PIK3CA* (252) but express mutant *TP53* (25–27). RG/C2s were cultured in Dulbecco’s Modified Eagles Medium (DMEM) (Life Technologies, Paisley, UK) and supplemented with 20% foetal bovine serum (FBS)(Life Technologies, Paisley, UK), L-glutamine (2mM)(Life Technologies, Paisley, UK), penicillin (100 units/ml) (Life Technologies, Paisley, UK), streptomycin (100 ug/ml) (Life Technologies, Paisley, UK) and insulin (0.2 units/ml) (Sigma-Aldrich, Poole, UK). Cells were mycoplasma tested (Mycoalert Plus mycoplasma detection kit; Lonza Group, Basal, Switzerland) and experiments performed within 10 passages. Aspirin (Sigma-Aldrich) was dissolved in fresh growth medium and diluted to form concentrations of 2mM and 4mM.

### Generation of proteomic data - SILAC approach

A stable isotope labelling with amino acids in cell culture (SILAC) approach was carried out on RG/C2 cells treated with 0mM, 2mM and 4mM aspirin for 24 hours. Control cells (0mM aspirin) were cultured with an L-arginine and L-lysine (light labelling), 2mM were cultured with ^2^H_4_-lysine and ^13^C_6_-arginine (medium labelling) and 4mM were cultured with ^15^N_2_ ^13^C_6_-lysine and ^15^N_4_ ^13^C_6_-arginine (heavy labelling) (Cambridge Isotope Laboratory, Massachusetts, United States). These methods were based on the SILAC-based mass spectrometry approach by Trinkle-Mulcahy et. al (2008) (28).

Cells were cultured with aspirin and the isotopes for 24 hours before extracting protein lysates. This experiment was carried out in duplicate. Lysates from the three conditions were pooled in a 1:1:1 ratio, separated by SDS-PAGE and then subjected to in-gel tryptic digestion. The resulting peptides were analysed by liquid chromatography mass spectrometry using an LTQ Orbitrap Velos mass spectrometer (Thermo Fisher Scientific, Waltham, Massachusetts, USA) and the mass spectral data analysed using Proteome Discoverer software v1.4 (Thermo). Details of SILAC labelling and proteomics have been previously published (29). To determine proteins whose expression is altered due to aspirin treatment, we applied a threshold of a 1.4 fold change between 4mM/control and 2mM/control, as suggested previously (30). Results were also limited to a variability of <100% and a peptide count of at least 2.

### Statistical analyses

#### Two-sample MR

To assess the effect of protein/mRNA expression of aspirin targets on risk of CRC, we used a two-sample MR approach. Firstly, SNPs were identified to proxy for protein/mRNA expression of the proteins shown to be altered in cell culture. Genetic association estimates with protein/mRNA expression levels (pQTLs/eQTLs) (sample 1) were integrated with genetic association estimates with CRC risk (sample 2).

#### Genetic predictors for protein and gene expression

Protein quantitative trait loci (pQTLs) were obtained from the INTERVAL study which comprises about 50,000 individuals within a randomised trial evaluating the effect of varying intervals between blood donations and how this affects outcomes such as quality of life (31). Relative protein measurements were taken using SOMAscan assays for 3,622 plasma proteins in a subset of 3,301 participants, randomly chosen. Genotyping and imputation (using a combined 1000 Genomes Phase 3-UK10K as the reference panel) of these individuals provided measures for 10,572,814 variants that passed quality control and were taken forward in a GWAS analysis to identify pQTLs for the measured proteins (details of quality control are mentioned elsewhere (19)). pQTLs identified represent a standard deviation (SD) change in protein expression (19).To adjust for multiple testing, a Bonferroni correction (0.05/10,572,814=4.72×10^−9^) was applied and pQTLs below this P-value threshold were used to proxy for protein expression in our analysis (32).

In the absence of a relevant pQTL for the protein of interest, an equivalent mRNA expression GWAS was used instead. Expression quantitative trait loci (eQLTs) were extracted from the eQTLGEN consortium consisting of 31,684 individuals from 37 datasets, of which 26,886 samples were from blood and 4798 from peripheral blood mononuclear cells (PBMCs). Due to the differing methods for genotyping between the studies, variants for each transcript ranged between 2,337-31,684 variants (20). For this reason, a Bonferroni correction threshold was adjusted depending on the number of variants measured for each transcript (0.05/number of variants) (32). eQTLs were standardized and meta-analysed through a Z-transformation, therefore eQTL effect sizes are reported as standard deviation (SD) changes (20).

In this analysis, both cis (within 1 Mb of the gene transcription start sit) and trans QTLs were used to proxy for expression. Once suitable pQTLs/eQTLs were identified, linkage disequilibiurm (LD) clumping at an R^2^ of 0.001 was carried out to remove SNPs that are inherited together and so that only the SNP most strongly associated with the mRNA/protein expression within a 10,000kb window was used.

#### Genetic association for colorectal cancer

Genetic association summary statistics for CRC, comprising 55,168 colorectal cancer cases and 65,160 controls, were obtained from the Colon Cancer Family Registry (CCFR), Colorectal Transdisciplinary (CORECT) and Genetics and Epidemiology of Colorectal Cancer (GECCO) consortia and UK Biobank (33–35). Quality control procedures have been described elsewhere (33). Ethics were approved by respective institutional review boards.

#### Evaluating the association of mRNA/protein expression on colorectal cancer

Analyses were carried out in R version 3.2.3 using the MR-Base TwoSampleMR R package (github.com/MRCIEU/TwoSampleMR) (36), which allows the formatting, harmonisation and analysis of summary statistics. The package reassigns alleles so that the effect allele has a positive association with the exposure and so represents an increase in protein/mRNA expression. In turn, allele harmonization ensures that the same allele (that predicts increased expression) is the effect allele in the outcome dataset as well. In the case of palindromic SNPs (represented by either A/T or G/C on both the forward and reverse alleles) these were also harmonized where possible based on allele frequencies. If allele frequencies for the effect allele and the other allele were similar, thus making harmonization difficult, these SNPs were dropped from the analysis (36).

Separate MR analyses were carried for cis and trans pQTLs as well as cis and trans eQTLs. For proteins with just one pQTL or eQTL, Wald ratios (SNP-outcome estimate ÷ SNP-exposure estimate) were calculated to give a causal estimate for risk of CRC per SD increase in mRNA/protein expression. Where more than one QTL was available as a proxy for the exposure (mRNA/protein levels), a weighted mean of the ratio estimates weighted by the inverse variance of the ratio estimates (inverse-variance weighted (IVW) method) was used (37).

When one genetic variant used to proxy for an exposure is invalid e.g. due to horizontal pleiotropy (where a genetic variant affects the outcome through an alternative exposure/pathway of interest) (17), then the estimator from the IVW method becomes biased (38). As a sensitivity analysis, alternative MR methods were used when more than 2 SNPs were available as instruments for mRNA/protein expression (MR Egger, simple mode, weighted mode, and weighted median) (36,39,40). Unlike the IVW method, the MR Egger method is not constrained to pass through an effect size of 0, thereby allowing the assessment of horizontal pleiotropy through the y intercept. (38,41). The weighted median approach is useful as it allows a consistent estimate even if 50% of the SNPs proxying protein/mRNA expression are invalid instruments (40) and the mode estimate also provides a consistent causal effect estimate even if the majority of the instruments are invalid, as the estimate depends on the largest number of similar instruments (39).

## Results

### Mendelian randomization of gene/protein expression and risk of colorectal cancer identified in aspirin treated human adenoma cells

In order to investigate the early changes that could reduce cancer risk, we investigated the proteome of aspirin treated adenoma derived cells to identify new targets of aspirin that may alter the risk of CRC by combining these proteomic results with an MR analysis. After applying a filtering threshold based on fold change and variability in expression, we identified 125 proteins whose expression appeared to be regulated by aspirin treatment (Figure 1) (S1 Table), although 5 were uncharacterised from mass spectrometry and therefore excluded from the analysis.

**Figure 1.**
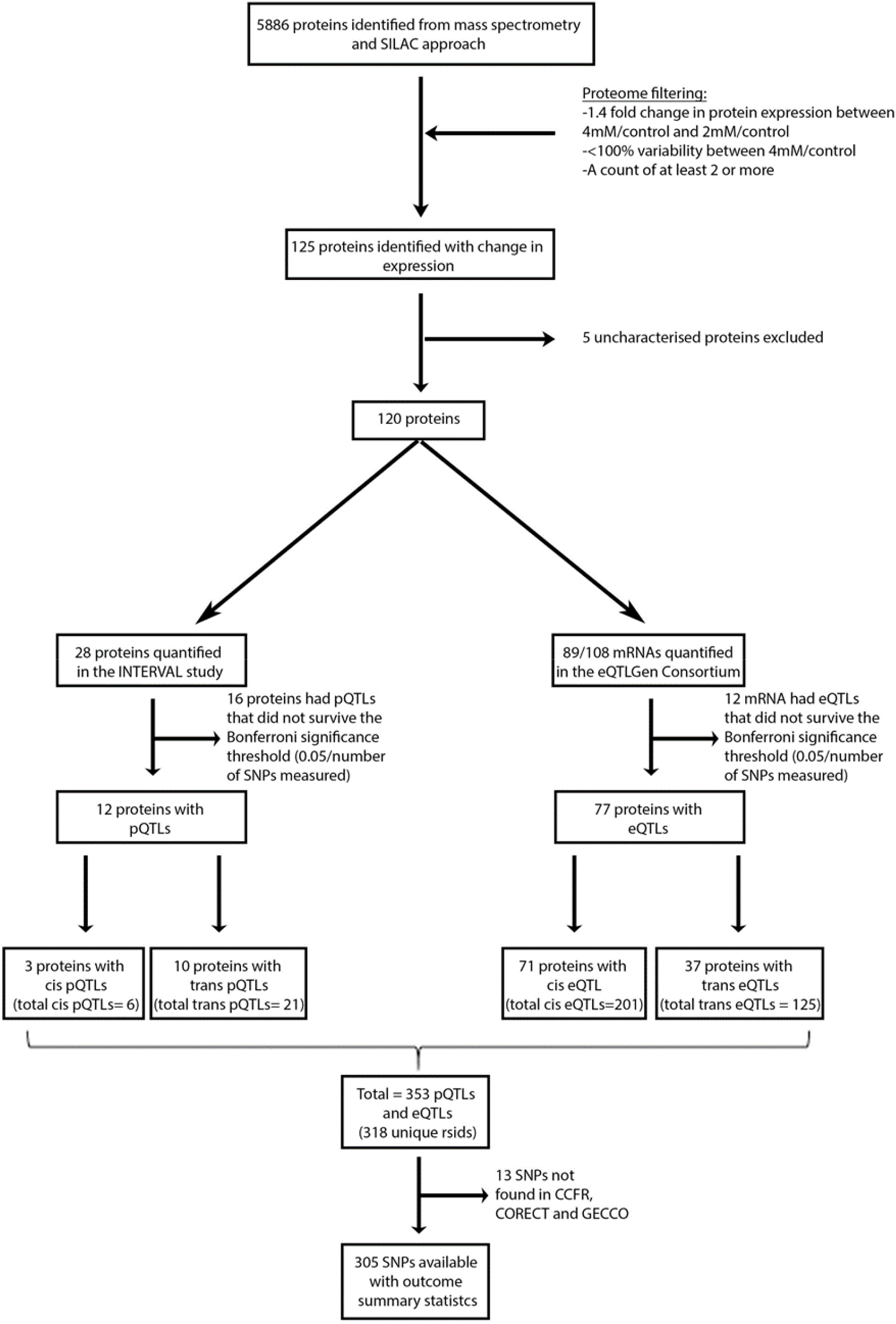
Flow diagram of SNP selection. 5886 proteins were identified using the SILAC proteomic approach. After applying a threshold, 125 proteins appear to be regulated by aspirin treatment, of which 5 were uncharacterised proteins and were therefore excluded from the analysis. In total, 12 proteins and 77 mRNAs had been quantified and had pQTLs/eQTLs below the Bonferroni significance threshold. Overall, summary statistics for 353 pQTLs and eQTLs were available, of which summary statistics for 305 of the SNPs was also present in the CCFR, CORECT and GECCO consortia.

Of the 120 proteins, expression of 28 proteins was measured in the INTERVAL study, of which 12 proteins had pQTLs that were below the Bonferroni significance threshold (0.05/10,572,814 = 4.73 ×10^−9^). From these 12 proteins, cis pQTLs were available for 3 proteins and trans pQTLs for 10 proteins (S2 Table). In the absence of available pQTLs, eQTLs for the transcripts of the identified proteins were used instead. Of the 108 proteins with no pQTLs available, expression of 89 mRNAs were measured in the eQTLGen consortium, of which 77 proteins had eQTLs that were below the Bonferroni significance threshold. From these 77 proteins, cis eQTLs were available for 71 proteins and trans eQTLs were available for 37 proteins (S3 Table). In total, there were 318 unique SNPs proxying for protein and mRNA expression, of which outcome summary statistics were available for 305 SNPs to test for association between 99 mRNA/proteins against risk of CRC.

Two-sample MR analysis using the Wald ratio or IVW method was conducted to test the effect of increased mRNA/protein expression on the risk of CRC incidence using cis and trans pQTLs (S4 Table) as well as cis and trans eQTLs (S5 Table). In total, 99 proteins were tested for association with CRC incidence. To correct for multiple testing, a Bonferroni adjusted threshold of significance was applied (0.05/99= 5.05×10^−4^) but we also considered associations of a nominal significance (P value<0.05). Overall, 1 protein with cis eQTLs and 2 with trans eQTLs were associated with CRC incidence at P< 5.05×10^−4^ and a further 3 proteins with cis eQTLs, 1 with a trans eQTL and 1 instrumented by a trans pQTL were associated with CRC incidence at a P value < 0.05.

Increased mRNA expression of Human Leukocyte Antigen A (*HLA-A*) and mini chromosome maintenance 6 (*MCM6*) instrumented by cis eQTLs was found to be associated with an increased risk of CRC incidence (OR 1.28, 95% CI:1.04-1.58, P value: 0.02 and OR 1.08, 95% CI: 1.03-1.13, P value: 9.23×10^−4^ per SD increase in mRNA expression, respectively). An SD increase in mRNA expression of fatty acid desaturase 2 (*FADS2*) and DNA polymerase delta subunit 2 (*POLD2*) instrumented by cis eQTLs was associated with a decrease in risk of CRC incidence (OR 0.94, 95% CI: 0.90-0.97, P value: 2.50×10^−4^ and OR 0.84, 95% CI: 0.75-0.94, P value: 1.17×10^−3^, respectively) (Figure 2, Table 1). For *FADS2* and *POLD2*, results were consistent using other MR methods (weighted median, weighted mode and simple mode) and the MR Egger test shows no evidence of pleiotropy (S6 Table, Supplementary Figure 1). From the cis eQTL analysis, only results for *FADS2* survived the Bonferroni significance threshold.

**Table 1.**
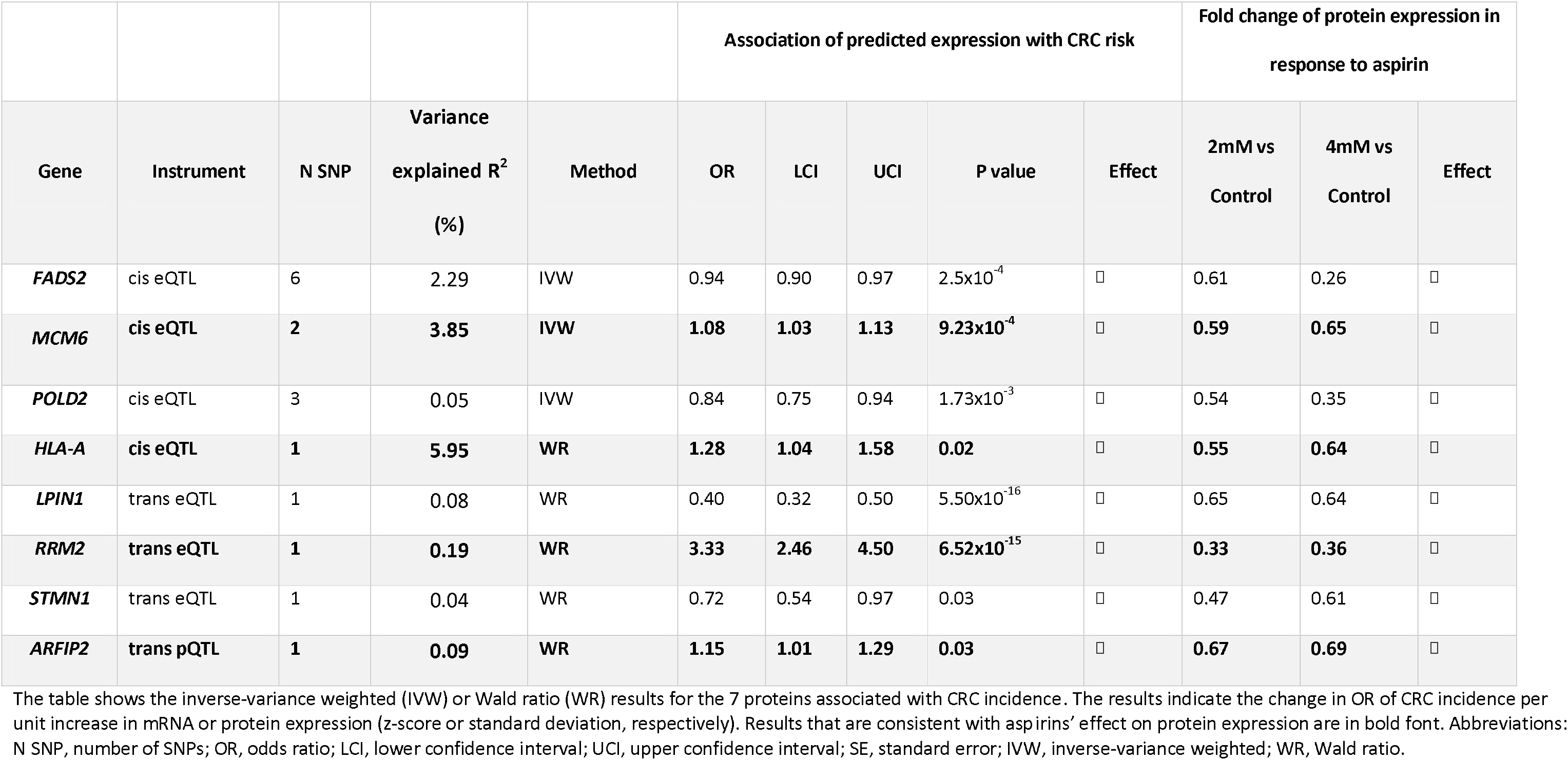
MR results of the 8 proteins associated with CRC incidence

| Gene | Instrument | N SNP | Variance<br>explained R<sup>2</sup><br>(%) | Method | Association of predicted expression with CRC risk |  |  |  |  | Fold change of protein expression in<br>response to aspirin |  |  |
| --- | --- | --- | --- | --- | --- | --- | --- | --- | --- | --- | --- | --- |
|  |  |  |  |  | OR | LCI | UCI | P value | Effect | 2mM vs<br>Control | 4mM vs<br>Control | Effect |
| *FADS2* | cis eQTL | 6 | 2.29 | IVW | 0.94 | 0.90 | 0.97 | 2.5x10<sup>-4</sup> | □ | 0.61 | 0.26 | □ |
| *MCM6* | cis eQTL | 2 | 3.85 | IVW | 1.08 | 1.03 | 1.13 | 9.23x10<sup>-4</sup> | □ | 0.59 | 0.65 | □ |
| *POLD2* | cis eQTL | 3 | 0.05 | IVW | 0.84 | 0.75 | 0.94 | 1.73x10<sup>-3</sup> | □ | 0.54 | 0.35 | □ |
| *HLA-A* | cis eQTL | 1 | 5.95 | WR | 1.28 | 1.04 | 1.58 | 0.02 | □ | 0.55 | 0.64 | □ |
| *LPIN1* | trans eQTL | 1 | 0.08 | WR | 0.40 | 0.32 | 0.50 | 5.50x10<sup>-16</sup> | □ | 0.65 | 0.64 | □ |
| *RRM2* | trans eQTL | 1 | 0.19 | WR | 3.33 | 2.46 | 4.50 | 6.52x10<sup>-15</sup> | □ | 0.33 | 0.36 | □ |
| *STMN1* | trans eQTL | 1 | 0.04 | WR | 0.72 | 0.54 | 0.97 | 0.03 | □ | 0.47 | 0.61 | □ |
| *ARFIP2* | trans pQTL | 1 | 0.09 | WR | 1.15 | 1.01 | 1.29 | 0.03 | □ | 0.67 | 0.69 | □ |

4 The table shows the inverse-variance weighted (IVW) or Wald ratio (WR) results for the 7 proteins associated with CRC incidence. The results indicate the change in OR of CRC incidence per 5 unit increase in mRNA or protein expression (z-score or standard deviation, respectively). Results that are consistent with aspirins' effect on protein expression are in bold font. Abbreviations: 6 N SNP, number of SNPs; OR, odds ratio; LCI, lower confidence interval; UCI, upper confidence interval; SE, standard error; IVW, inverse-variance weighted; WR, Wald ratio.

**Figure 2.**
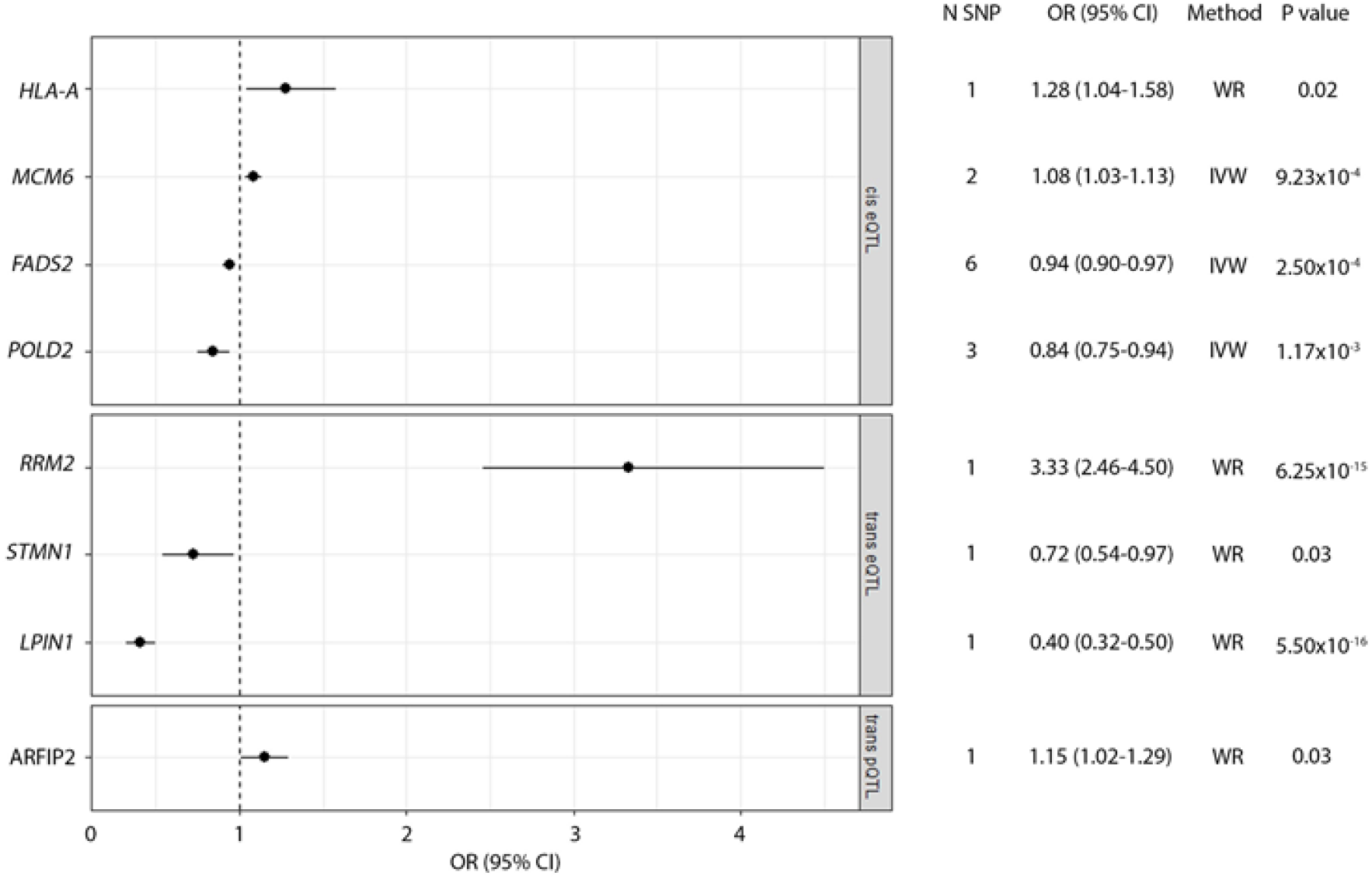
Forest plot of mRNA/protein associations with CRC incidence at a P value of <0.05. The upper box presents results using cis eQTLs, followed by trans eQTLs and finally trans pQTLs. Each dot on the plot represents the change in OR of CRC incidence per SD increase in mRNA/protein expression and the horizontal lines either side of the dot represent the 95% confidence intervals. The dotted line represents a null association between expression and cancer incidence. The number of SNPs used as instruments as well as the OR, the method and P value of association are also reported. Abbreviations: N SNP, number of SNPs; OR, odds ratio; CI, confidence intervals; IVW, inverse-variance weighted; WR, Wald ratio.

Proteins instrumented by trans eQTLs include ribonucleoside-diphosphate reductase subunit M2 (*RRM2*), stathmin-1 (*STMN1*) and lipin 1 (*LPIN1*). An increase in *RRM2* was estimated to increase the risk of cancer incidence (OR 3.33, 95% CI: 2.46-4.50, P value: 6.25×10^−15^ per SD increase in mRNA expression) whereas an increase in *STMN1* and *LPIN1* was associated with decreases in the risk of CRC incidence (OR 0.72, 95% CI: 0.54-0.97, P value: 0.03 and OR 0.40, 95% CI: 0.32-0.50, P value: 5.50×10^−16^ per SD increase in mRNA expression, respectively). From the trans eQTL analysis, results for *RRM2* and *LPIN1* both survived the Bonferroni significance threshold.

For proteins instrumented by pQTLs, ADP ribosylation factor interacting protein 2 (ARFIP2) proxied using a trans pQTL conferred an increased risk of CRC incidence (OR 1.15, 95% CI: 1.01-1.29, P value: 0.03 per SD increase in protein expression).

Overall, the directions of effects between *HLA-A*, *MCM6*, *RRM2* and *ARFIP2* and CRC risk obtained from our MR analysis concur with those anticipated given the protective role of aspirin on CRC and the effect of aspirin treatment on expression of these proteins. Aspirin reduces the protein expression of HLA-A, MCM6, RRM2 and ARFIP2 (fold change in protein expression with 4mM aspirin treatment compared to control: 0.55, 0.65, 0.36 and 0.69, respectively, Table 1) and aspirin intake is associated with a decreased risk of CRC (2–4). Our MR analysis shows that increased expression of these proteins is associated with an increased risk of CRC incidence. Taken together, our results indicate that a possible mechanism through which aspirin decreases the risk of CRC incidence is through the downregulation of HLA-A, MCM6, RRM2 and ARFIP2. The direction of effect was less consistent for the other 4 proteins (FADS2, POLD2, STMN1 and LPIN1) showing opposite results to what we would expect based on the proteomic results (Table 1).

## Discussion

Evidence for the use of aspirin in the prevention of CRC is increasing (2–5). However, the mechanism through which it functions is still not fully understood. By combining both a proteomic-based approach as well as an MR analysis, our results provide mechanistic insights into how aspirin could decrease the risk of CRC.

Using a SILAC-based proteomics approach, 120 proteins appear to be regulated at 24 hours by 4mM and 2mM aspirin treatment. Genetic variants (pQTLs and eQTLs) were identified and used to proxy for protein and mRNA expression levels of the identified proteins to test for evidence of a causal effect on CRC incidence. When no pQTL was available for a protein, eQTLs were used instead.

Overall, 4 cis eQTLs, 3 trans eQTLs and 1 trans pQTL were associated with cancer incidence at a P value < 0.05. Increased expression of *HLA-A* and *MCM6* proxied by cis eQTLs were associated with an increase in the risk of CRC incidence and an increase in *RRM2* and ARFIP2 (proxied by a trans eQTL and trans pQTL, respectively) also conferred an increased risk. Therefore, suppressing the expression of these four proteins could decrease the risk of CRC. As the proteomic results showed that aspirin treatment decreases the expression of these proteins, this could be a potential mechanism by which aspirin reduces the risk of CRC. However, only results for *RRM2* survive the Bonferroni significance threshold, indicating that further studies are required to verify these results.

The proteins MCM6 and RRM2 are both involved in repair of DNA damage. MCM6 is part of a helicase complex involved in unwinding DNA and is involved in repair of double stranded breaks (DSBs) in homologous recombination through interaction with RAD51. This interaction is required for chromatin localisation and formation of foci for DNA damage recovery (42). Likewise, RRM2 is part of a protein complex called ribonucleotide reductase which catalyses the biosynthesis of dNTPs and is therefore required for DNA replication and damage repair (43).

Cancer cells commonly lose the DNA damage response, which results in the accumulation of mutations that may be oncogenic (44). Because of this, tumour cells end up relying on a reduced number of repair pathways and are therefore more sensitive to inhibition of DNA damage repair pathways when compared to normal cells which have full capability of DNA repair (45). Drugs that target these other pathways have been shown to selectively kill the cancer cells which is known as synthetic lethality (46,47). It may be that by reducing the expression of DNA repair proteins, which combined with DNA damage response proteins that are already mutated during tumour progression, aspirin can induce cell death in the developing tumour cells reducing the risk of developing cancer.

The MR results for the proteins ARFIP2 and HLA-A also concur with our SILAC proteomic results. ARFIP2 is a protein previously shown to play a role in membrane ruffling and actin polymerization, therefore regulating the actin cytoskeleton (48). The remodelling of the actin cytoskeleton is known to be involved in cancer metastasis (49). This is of particular interest as aspirin reduces the odds of colorectal adenocarcinoma metastasis by 64% (OR:0.36 (95% CI: 0.18-0.74)) (50) and this may be through the reduction in ARFIP2 expression. With regards to HLA-A expression and cancer risk, results from a cohort study showed that aspirin was more chemopreventative in tumours that expressed HLA class I antigen (which includes HLA-A, HLA-B and HLA-C) (rate ratio (RR) 0.53, 95% CI: 0.38-0.74) and this association was no longer apparent in tumours that lacked expression of this protein (15). Our MR analysis showed that an increase in *HLA-A* was associated with increased CRC risk, and that aspirin may reduce this risk through a reduction in HLA-A expression, however further investigation is required before any conclusions can be drawn.

Our MR analysis results also showed that increased mRNA expression of *FADS2*, *POLD2*, *LPIN1* and *STMN1* all decreased the risk of CRC, indicating that decreased expression increases the risk of cancer. Our proteomic results showed that aspirin decreases the expression of these proteins and aspirin is known to decrease cancer risk. The exact meaning behind the inconsistencies in direction of effect is unclear but may be related to the dosage used in this study. A randomized trial of aspirin to prevent adenomas showed that lower doses reduced adenoma risk more than higher doses, suggesting that lower doses of aspirin may affect mRNA/protein expression differently than higher doses (51,52). Furthermore, the genetic instruments used to proxy for *POLD2*, *LPIN1* and *STMN1* explain little of the variance in mRNA expression (0.05, 0.08 and 0.04%, respectively) indicating that SNPS that explain more of the variance are required before any conclusions can be made.

Further limitations also exist in our analysis. Firstly, the exact correlation between eQTLs and pQTLs has not been fully determined. Secondly, it is difficult to interpret results using trans eQTLs and pQTLs without clear confirmation that these SNPs directly influence the gene/protein expression. It may be that they indirectly influence expression, for example, trans eQTLs may regulate gene expression by affecting expression of a nearby cis gene which is in fact a transcription factor that is regulating the expression of the trans gene (53). Thirdly, both the pQTL and eQTL associations were carried out using blood samples or PBMCs (19,20), therefore these SNPs estimate changes in gene and protein expression in circulating immune cells only. As found by the Genotype-Tissue Expression (GTEx) study, cis eQTLs are either shared across tissues or are specific to a small number of tissues (54). Therefore, the use of these eQTLs and pQTLs measured in the blood may not be fully suitable as proxies for mRNA and protein expression in the epithelium of the colon and rectum. Furthermore, the units for the eQTLs and pQTLs represent SD changes in expression, making interpretation of the results difficult. However, we can interpret the direction of effect as well as the statistical significance of the association (P values) for these analyses. Moreover, pQTLs and eQTLs could not be identified for 20 of the proteins found to be regulated by aspirin in our proteomic approach, therefore we could not test the association of their expression with CRC risk. Finally, apart from the association of *FADS2* with CRC incidence, the other associations proxied by cis eQTLs found by our study are not below the Bonferroni threshold of significance (P value ≤ 4.63×10^−4^).

MR is commonly used to proxy for a drug’s effect on risk of various outcomes after identification of its target. Genetic variants that predict lower function of 3-hydroxy-3-methylglutaryl coenzyme A (HMG-CoA) reductase are commonly used to investigate the effect of lowering LDL cholesterol via the use of statins on outcomes such as ovarian cancer, Alzheimer’s disease or coronary heart disease (55–57). These studies involve investigation of a drug’s effect via a known target on an outcome. However, this approach would be difficult to apply in the case of drugs with pleiotropic targets such as aspirin. Therefore, in order to identify all possible targets of aspirin, a proteomic approach was firstly applied and targets that may affect risk of cancer were identified through using MR. To our knowledge, this is the first study that combines basic science and MR to generate hypotheses of a drug’s mechanism of action in cancer.

Further experiments need to be conducted to confirm the effect of aspirin on gene and protein expression and the consequent effect this may have on hypothesised pathways such as DNA repair before definitive conclusions can be made. However, the potential of this unbiased approach to gain mechanistic insight is clear, allowing hypothesis driven research will better inform the clinical use of aspirin for the prevention of CRC.

## Acknowledgements

ASTERISK: We are very grateful to Dr. Bruno Buecher without whom this project would not have existed. We also thank all those who agreed to participate in this study, including the patients and the healthy control persons, as well as all the physicians, technicians and students.

CLUE: We appreciate the continued efforts of the staff members at the Johns Hopkins George W. Comstock Center for Public Health Research and Prevention in the conduct of the CLUE II study. We thank the participants in CLUE. Cancer incidence data for CLUE were provided by the Maryland Cancer Registry, Center for Cancer Surveillance and Control, Maryland Department of Health, 201 W. Preston Street, Room 400, Baltimore, MD 21201, http://phpa.dhmh.maryland.gov/cancer, 410-767-4055. We acknowledge the State of Maryland, the Maryland Cigarette Restitution Fund, and the National Program of Cancer Registries of the Centers for Disease Control and Prevention for the funds that support the collection and availability of the cancer registry data.

COLON and NQplus: the authors would like to thank the COLON and NQplus investigators at Wageningen University & Research and the involved clinicians in the participating hospitals.

CORSA: We kindly thank all those who contributed to the screening project Burgenland against CRC. Furthermore, we are grateful to Doris Mejri and Monika Hunjadi for laboratory assistance.

CPS-II: The authors thank the CPS-II participants and Study Management Group for their invaluable contributions to this research. The authors would also like to acknowledge the contribution to this study from central cancer registries supported through the Centers for Disease Control and Prevention National Program of Cancer Registries, and cancer registries supported by the National Cancer Institute Surveillance Epidemiology and End Results program.

Czech Republic CCS: We are thankful to all clinicians in major hospitals in the Czech Republic, without whom the study would not be practicable. We are also sincerely grateful to all patients participating in this study.

DACHS: We thank all participants and cooperating clinicians, and Ute Handte-Daub, Utz Benscheid, Muhabbet Celik and Ursula Eilber for excellent technical assistance.

EDRN: We acknowledge all the following contributors to the development of the resource: University of Pittsburgh School of Medicine, Department of Gastroenterology, Hepatology and Nutrition: Lynda Dzubinski; University of Pittsburgh School of Medicine, Department of Pathology: Michelle Bisceglia; and University of Pittsburgh School of Medicine, Department of Biomedical Informatics.

EPIC: Where authors are identified as personnel of the International Agency for Research on Cancer/World Health Organization, the authors alone are responsible for the views expressed in this article and they do not necessarily represent the decisions, policy or views of the International Agency for Research on Cancer/World Health Organization.

EPICOLON: We are sincerely grateful to all patients participating in this study who were recruited as part of the EPICOLON project. We acknowledge the Spanish National DNA Bank, Biobank of Hospital Clínic–IDIBAPS and Biobanco Vasco for the availability of the samples. The work was carried out (in part) at the Esther Koplowitz Centre, Barcelona.

Harvard cohorts (HPFS, NHS, PHS): The study protocol was approved by the institutional review boards of the Brigham and Women’s Hospital and Harvard T.H. Chan School of Public Health, and those of participating registries as required. We would like to thank the participants and staff of the HPFS, NHS and PHS for their valuable contributions as well as the following state cancer registries for their help: AL, AZ, AR, CA, CO, CT, DE, FL, GA, ID, IL, IN, IA, KY, LA, ME, MD, MA, MI, NE, NH, NJ, NY, NC, ND, OH, OK, OR, PA, RI, SC, TN, TX, VA, WA, WY. The authors assume full responsibility for analyses and interpretation of these data.

Kentucky: We would like to acknowledge the staff at the Kentucky Cancer Registry.

LCCS: We acknowledge the contributions of Jennifer Barrett, Robin Waxman, Gillian Smith and Emma Northwood in conducting this study.

NCCCS I & II: We would like to thank the study participants, and the NC Colorectal Cancer Study staff.

NSHDS investigators thank the Biobank Research Unit at Umeå University, the Västerbotten Intervention Programme, the Northern Sweden MONICA study and Region Västerbotten for providing data and samples and acknowledge the contribution from Biobank Sweden, supported by the Swedish Research Council (VR 2017-00650).

PLCO: The authors thank the PLCO Cancer Screening Trial screening center investigators and the staff from Information Management Services Inc and Westat Inc. Most importantly, we thank the study participants for their contributions that made this study possible.

SCCFR: The authors would like to thank the study participants and staff of the Hormones and Colon Cancer and Seattle Cancer Family Registry studies (CORE Studies).

SEARCH: We thank the SEARCH team.

SELECT: We thank the research and clinical staff at the sites that participated on SELECT study, without whom the trial would not have been successful. We are also grateful to the 35,533 dedicated men who participated in SELECT.

WHI: The authors thank the WHI investigators and staff for their dedication, and the study participants for making the program possible. A full listing of WHI investigators can be found at: http://www.whi.org/researchers/Documents%20%20Write%20a%20Paper/WHI%20Investigator%20Short%20List.pdf

