## Supplementary Figure 1 for "A combined proteomics and Mendelian randomization approach to investigate the effects of aspirin-targeted proteins on colorectal cancer"

Supplementary Figure 1- Summary plots for FADS2 and POLD2

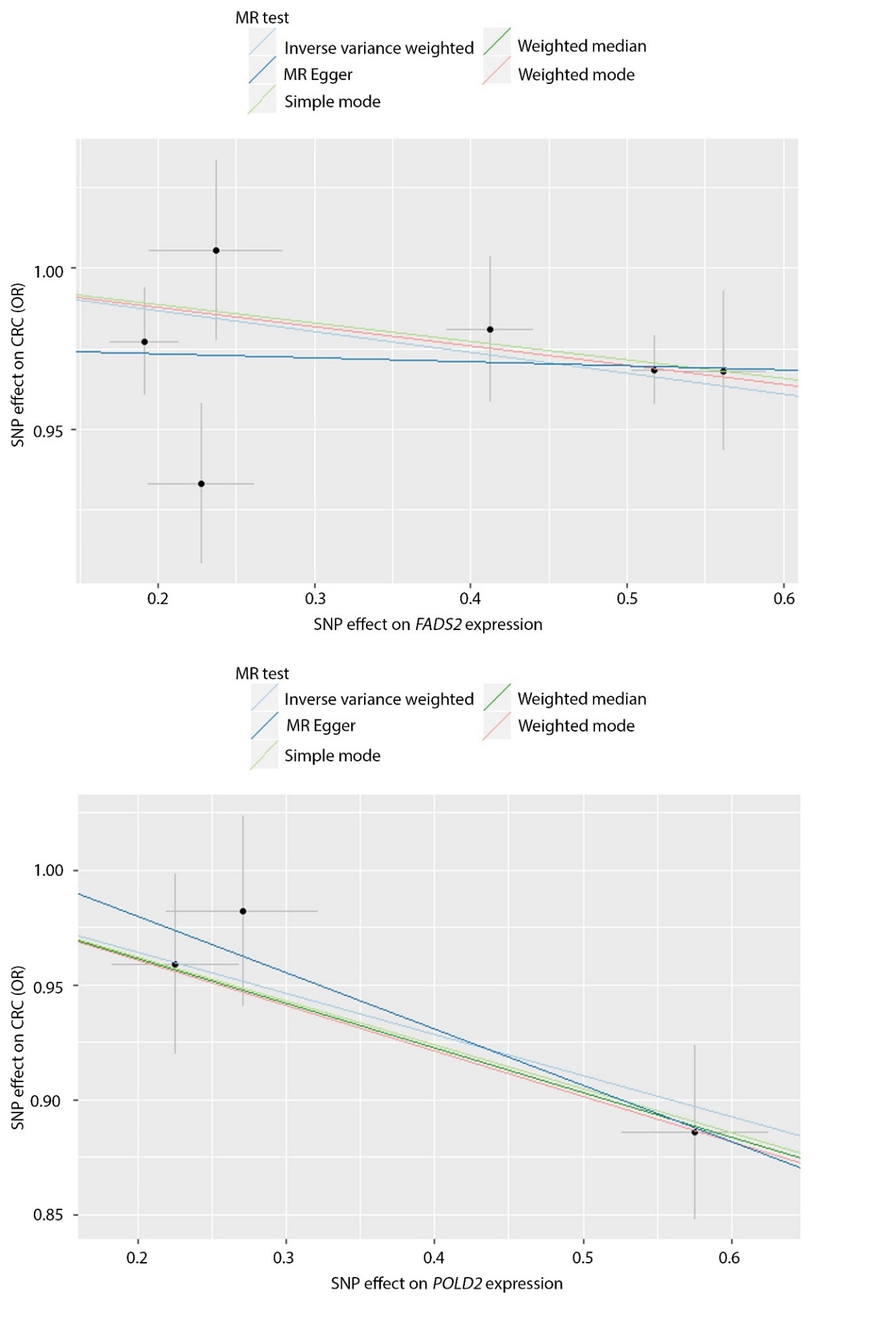

Scatter plot showing the effect of increasing *FADS2* and *POLD2* expression on risk of CRC. The black dots represent each SNP and the lines either side parallel to the x-axis and y-axis represent the 95% confidence intervals for SNP effect on gene expression and SNP effect on cancer risk, respectively. Five methods in total were used to summarise the results: inverse variance weighted (light blue), MR Egger (dark blue), Simple mode (light green), Weighted median (dark green) and weighted mode (light pink).
